# Stimulated oxygen transport in tissue by magnetic needle acupuncture

**DOI:** 10.1101/2021.02.01.426016

**Authors:** Zhu Liu, Chenyu Wen, Shi-Li Zhang

## Abstract

**Aim:** To understand physiology of magnetic needle acupuncture by investigating O_2_ transport in tissue during needle intervention.

**Methods:** O_2_ transport in tissue is modeled by utilizing COMSOL with magnetic needle inserted into muscle tissue in a 2D porous media. The damaged tissue has been mimicked by an extracted tissue block with 1^st^ order O_2_ consumption rate. The convection-diffusion O_2_ transport in the damaged tissue has been further evaluated by varying magnetic flux density *B*_0_ of the needle (0-1 T), myoglobin concentration (0-1 mM), O_2_ tension (5-100 Torr), O_2_ consumption rates, tissue permeability (10^−12^-10^−6^ m^2^) and porosity (0.1-0.9).

**Results:** 1) Active O_2_ transport carried by interstitial flow is enhanced with the intervention of a magnetic needle by generating a high gradient magnetic field around the tip, which exerts a strong force (10^4^ N/m^3^) on the diamagnetic interstitial fluid to accelerate the flow. 2) This interstitial flow can reach 30 μm/s at *B*_0_ = 1T and strongly correlates to the needle *B*_0_ and tissue permeability. 3) The needle stirs the interstitial flow can pump O_2_ flux by 1-2 orders of magnitude compared to that without magnetic field. 4) The enhancement of active O_2_ transport by magnetic needle is site-specific to the tissue in the vicinity of the tip. This enhancement is more effective in edema condition with a high tissue permeability (>10^−9^ m^2^).

**Conclusions:** The dramatic enhancement of O_2_ transport to restore the O_2_ mitochondria metabolism for dysfunctional muscle tissues is the fundamental physiological mechanism of magnetic needle acupuncture.

## 1. Introduction

Chinese acupuncture has been used to treat different diseases for more than 2000 years. During the acupuncture process, the needle is inserted into the muscle tissue at certain acupoints, followed by twirling the needle to generate “de-qi” feeling of distension, heaviness, numbness, soreness, etc. ^1^ Typically, local microcirculation with improved blood perfuse ^2–4^ has also been observed after the intervention of the acupuncture needle. By time, different techniques have been fused into the acupuncture therapy such as electroacupuncture^5,6^, magnetic acupuncture ^7^, laser acupuncture ^8^, microwave acupuncture ^9^, etc. The acupuncture needles are standardized with respect to shape and material. This far, however, acupuncture has yet to reach beyond being treated as an alternative therapy for chronic pain relief ^10^, systemic inflammation ^11^, depression ^12^, anesthesia ^13^, etc., though progressively gaining credibility as a primary or adjuvant therapy.

However, acupuncture therapy has persistently been questioned as a placebo ^14,15^, in spite of robustly documented clinical recovering cases and patients. Its major controversy is the physiological mechanisms, based on the primitive meridian channel theory of the traditional Chinese medicine system ^16^, that lack full support in the framework of modern medical and physiological theorems. Moreover, the acupuncture technique has been dramatically advanced without the understanding of fundamental physiological processes, which has also made the mechanism of the acupuncture difficult to validate. As a result, suspicion persists on the acupuncture treatment.

In the past decades, different theories based on neuron excitation ^6,17^, microcirculation, mechanical signaling through connective tissue ^18^, and acupoint anatomy ^19^ have been proposed to account for the therapy effects of acupuncture, as well as the “de-qi” feeling during the needle intervention. The neuron excitation of thin A-δ and unmyelinated C-fibers ^17,20^ in the skin and muscle has been suggested as the mechanism for the “de-qi” feeling; it is triggered by the modulation of release of endogenous endorphins and neurotransmitters for the central nervous system upon modulating the transmission of pain signals resulting in ^21^. The automatic sympathetic neuron activation during the acupuncture intervention has also been correlated to the production of NO or vasoactive neuropeptides to modulate the vasodilation of the blood vessel to enhance the local blood volume ^22,23^. In the other perspective, a consistent effort was on finding the anatomy acupoint sites in the meridian channel, by correlating to facial trigger points ^24,25^, skin resistance ^26^, and interstitial flow ^19^. Among these attempted explanations, the neuron activation for both central nervous system and sympathetic nervous system is believed to be a clue to understand the acupuncture. However, several puzzles remain: why such a simple needle manipulation can generate similar nociceptive feelings but secret dissimilar hormones for different diseases; how the needle can find the specific neuron to generate the specific molecules; why there is “de-qi” feeling; etc. Moreover, the neuron excitation cannot be used for the laser acupuncture since there is no physical puncture through the skin and no excited neuron.

A different approach has been introduced based on the mitochondria metabolism related to the laser acupuncture inducing the local microcirculation. Since the local O_2_ metabolism plays a crucial role in cellular energetics and cell recovery ^8^, the mechanism of the laser acupuncture was attributed to the mitochondria absorption of the red visible and near infrared light thereby being a part of the cellular respiratory chain. However, this implication is constrained to the optical absorption of molecules, involved in the mitochondria’s O_2_ metabolism. It has nothing in common with the physical puncture of the traditional acupuncture. Unfortunately, there is no further physiological model relating the mechanism of the acupuncture (dry needle) to the local O_2_ metabolism.

Many diseases are correlated to different damages of the muscle tissue or reduced adenosine triphosphate (ATP) generation by O_2_ mitochondria metabolism. Typically, pain in muscle tissue is correlated to the hypoxia or ischemia condition of tissue ^27^ and impeachment of O_2_ metabolism. Consistently low local blood flow was found in the painful tissue with elevated muscle tension ^28,29^. The biopsy of the chronic myalgia shows abnormal muscle fibers with “ragged red” and “moth eaten” corresponding to the ischemia condition. More studies found that the pain region has impairment of capillaries, drop of ATP generation, and abnormality of mitochondria ^27,30,31^. Hence, the damaged tissue with reduced capillaries becomes the obstacle for O_2_ transport inside tissue and O_2_ mitochondria metabolism. Furthermore, the stiffness and reduced contraction muscle tissue impeach the lymph flow and worsen the microcirculation. It has been suggested that mitochondrial, microcirculatory, and/or metabolic changes in the trapezius muscle sensitize muscle nociceptors in patients ^27^. Therefore, to recover the tissue from pain and reduced O_2_ mitochondria condition, it is crucial to restore the O_2_ transport through the tissue, to repair the damage tissue, and to rehabilitate mitochondria metabolism.

Recently, the non-vessel interstitial flow has been proposed to explain the meridian channel theory ^19^. The meridian channel has been viewed to be located in the connective tissue with a high hydraulic conductance for the interstitial flow. In the meridian theory, the acupoint is the joint and connective point where the “qi” and “xue” is in and out from the channel.

In our previous study ^32^, a convection-diffusion model is built to describe the O_2_ transport in tissue by interstitial flow. This model elicits that the interstitial flow plays a crucial role in the O_2_ transport in tissue. Here, to understand the metabolism physiology of acupuncture, as a demonstration, a traditional magnetic needle has been used as the needle body and inserted into a two-dimensional (2D) porous media mimicking the muscle tissue. The simulation illustrates that the magnetic needle extorts a high gradient pressure on the interstitial fluid in tissue, due to an extremely high gradient magnetic field at tip that exerts a force on the water molecules of diamagnetic nature. This force drives a flow near the needle tip to stimulating active O_2_ transport, hence redistributes and pumps O_2_ into the tissue with a high hydraulic conductance. Therefore, the magnetic needle acupuncture can restore the O_2_ mitochondria metabolism for dysfunctional muscle tissues. The simulation is implemented on the COMSOL Multiphysics software package.

## 2. O_2_ transport in muscle tissue

The interstitial fluid flows through the interspace of muscle fibers. In muscle fiber, mitochondria is aggregated mainly near the subsarcolemmal region ^33^. The channels of transverse tubules, in sarcolemma of the muscle fiber, also penetrate into the fiber center and open up to the interstitial fluid ^34^. Hence, even the interstitial flow in the interspace can reach the center of the muscle fibers. Moreover, typically, water molecules can cross the cell membrane via aquaporins and its permeability depends on the pressure across the membrane for example the osmosis pressure caused by protein and ion concentration transmembrane gradient. Here, in our model, tissue permeability i.e., the specific hydraulic conductance includes the water permeability through both the interstitium and cell membrane of muscle fiber. Hence, the O_2_ transport in tissue through convection by the interstitial flow and the O_2_ diffusion by concentration gradient are considered during acupuncture retention.

### O_2_ transport inside muscle tissue

For muscle tissue, the concentration of O_2_ loaded myoglobin, *C*_Mb_O2_ can be described by the Hill equation at the reaction equilibrium state ^35^:

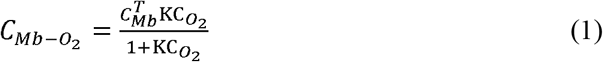

Here, *C*_Mb_^T^ is the total concentration of myoglobin (0-1 mM), including the O_2_-loaded sate and unloaded state. At low O_2_ concentration, myoglobin facilitates the O_2_ diffusion inside the tissue ^36^. *C*_O2_ = *αP*_O2_ is the concentration in capillary. *α* is the solubility coefficient of O_2_ in water (1.23×10^−3^ mol m^−3^ Torr^−1^) and *P*_O2_ is the O_2_ tension in capillary.

The passive O_2_ diffusive transport through tissue can be described by Fick’s first law:

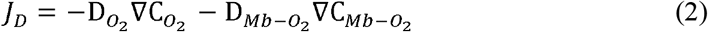

where, *D*_O2_ = 1.6×10^−9^ m^2^/s and *D*_Mb_=1 ×10^−10^ m^2^/s are the diffusion coefficient of O_2_ and myoglobin, respectively ^36^, assuming *D*_Mb_ = *D*_Mb_O2_ The convection contribution *J*_conv_ of the interstitial flow can be expressed as:

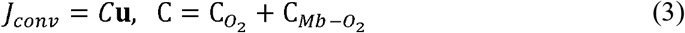

The total O_2_ flux is *J* = *J*_D_+*J*_conv_. The O_2_ input flux into the tissue is defined as the total flux *J* at *x* = 0, i.e., the O_2_ source. At steady state, *dC*_O2_/*dt* = 0, and the O_2_ flux distribution in the tissue per unit time per unit volume is described by Fick’s second law:

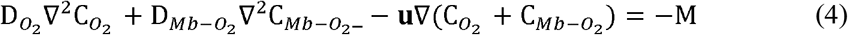

At rest, the O_2_ consumption rate *M* of mitochondria is around 5×10^−3^ mol·m^−3^·s^−1^ under the standard conditions (20 °C, 1 atm) ^37^. For impaired tissue, considering an impeached metabolism of mitochondria, the O_2_ consumption is assumed as a first order reaction and the ratio rate *M* depends on the O_2_ concentration (α*P*_O2_):

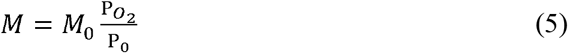

where, *M*_0_ is the O_2_ consumption rate (10^−3^ mol·m^-3^·s^−1^) under edema condition and *P*_0_ is the constant O_2_ tension in capillary.

### Interstitial flow in the muscle tissue

The interstitial flow rate correlates to the tissue permeability *κ* (m^2^) and porosity. For clarification, the tissue permeability here refers to the specific hydraulic conductance of tissue, not the permeability in electromagnetism. In this work, both tissue permeability and specific hydraulic conductance are used identically for convenience. The interstitial flow is described using a porous media with the Birkman equation ^38^:

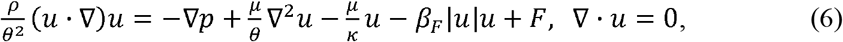

where, *θ* is the volume porosity of the interstitial region, factor *β*_F_ is the Forchheimer drag factor with the unit of kg·m^−4^, *μ* is the dynamic viscosity of water in 1×10^−3^ Pas, and ***F*** is the external body force on the fluid.

### Body force exerted by magnetic needle

A magnetic needle can provide a high gradient agnetic field, which generates a strong force on paramagnetic and diamagnetic molecules ^39 40^. At the tip, the magnetic flux density decays rapidly, rendering a high magnetic field gradient, which extorts a large body force on the diamagnetic fluid ^41,42^ and enhances O_2_ dissolution rate in water ^39^. In addition, this magnetic force can also enhance the paramagnetic ion transport ^43,44^ for the electrodeposition and ion separation. The gradient of the magnetic field at the tip can reach more than 10^3^ T/m and can be used to assist drug delivery ^40^.

The use of magnetic needle has a long history in the traditional Chinese acupuncture treatment. Therefore, the magnetic needle is a demonstration to understand the physiological effects of acupuncture on the impact of interstitial flow. The magnetic flux density *B*_0_ of the needle is determined by its volume magnetization (***M***_v_). For a fully magnetized needle ^40^, the following holds:

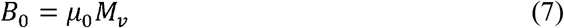

where, *μ*_0_ is the vacuum magnetic permittivity, 4π×10^−7^ H/m. ***M***_v_ depends on material type and magnetization process. For soft magnetic materials, the magnetic flux density after magnetized depends on the residue of induction and its shape anisotropy. *B*_0_ can reach 1-2 T in a magnetic needle made by iron after magnetization ^40^.

For a paramagnetic or diamagnetic fluid, the volume magnetic moment (emu/m^3^) in the magnetic field ***H*** generated by the magnetic needle can be expressed as:

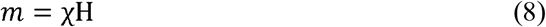

where, *χ* is the susceptibility of the fluid. The interstitial fluid of the muscle tissue is composited by water molecules, proteins and ions. If ignoring the magnetic properties of ions, *χ* relates to the components of the fluid via:

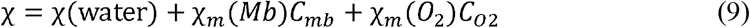

where, *χ_m_*(Mb) and *χ_m_*(O_2_) are the molar susceptibility of myoglobin and O_2_, respectively. Myoglobin without O_2_ loaded is a paramagnetic molecule with 5.48-5.8 *μ*_B_^45^, and O_2_ loaded myoglobin is a diamagnetic molecule. The susceptibility of water molecules is −9×10^−6^. Hence, the body force of the fluid in magnetic field ***H***, can be expressed as ^46^:

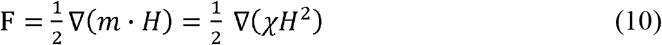

where, the factor 1/2 is due to the volume magnetic moment depending on ***H***.

### COMSOL simulation

The magnetic acupuncture is simulated by considering a needle inserted into a 2D porous media. In the simulation, the Nernst-Planck equation (NPE) is adopted to govern the dynamics of O_2_ and myoglobin, achieved by the transport of diluted species module. The Maxwell equation (ME) is used to describe the magnetic field distribution through the magnetic field module. The Navier-Stocks equation (NSE) is included to dictate the movement of the interstitial flow, mainly water, realized by the laminar flow module. These three modules are fully coupled. A body force acting on water of the interstitial flow is introduced by the gradient of magnetic field predicted by ME with equation (10). The interstitial flow velocity determined by NSE is fed to NPE for the convective transport. In the entire region governed by the NPE, the concentrations of O_2_, myoglobin, and O_2_ loaded myoglobin are connected by a reversible reaction at equilibrium, with the forward and reverse reaction rate constants *k*_1_ and *k*_2_ (k_1_ k_2_=6.9× 10^3^ m^3^ mol^−1^)

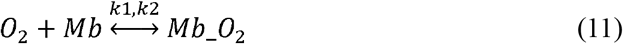

Thus, at equilibrium state, the concentrations follow equation (1). In addition, Birkman’s law of porous media is introduced in the laminar flow module.

The needle is 3.5 mm long with a diameter of 0.2 mm, and the tip has a radius of 0.5 mm, shown as domain ③ in **Fig. 1**. The magnetic field and interstitial flow distribution are calculated by following ME and NSE in the entire domain of 4 mm × 6.5 mm, i.e., domain ①+②+③ in **Fig. 1**. To monitor the O_2_ transport inside the muscle tissue, a tissue block ① of 1 mm × 3.8 mm right next (0.1 mm away) to the needle tip is extracted, illustrated in **Fig 1**. Side AB is set as the inflow while the other three sides, BC, CD, and DA, are set as the outflow in the laminar flow (NSE). For the O_2_ transport process (NPE), a constant *P*_O2_ = *P*_0_ is set at side EF, and the other 3 sides, i.e., FG, GH, and HE, are controlled by flow rates with ∇·*C* = 0. Before needle insertion into the tissue, the interstitial flow is set as stasis with ***u*** = 0. The tissue permeability is set as ***κ***=10^−8^ m^2^ under the damage edema condition. The temperature of tissue is at 37 °C. All parameters are list in **Table 1**.

**Fig. 1.**
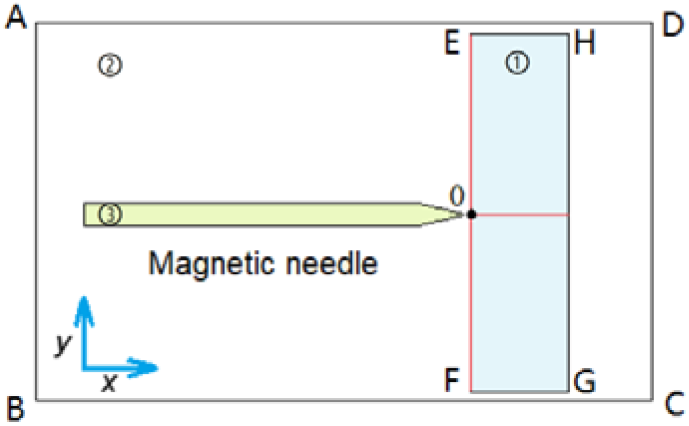
Definition of simulation domains. The magnetic field distribution of the needle ③ is calculated in the entire domain ①+②+③. The interstitial flow is calculated in domain ①+②, and the transport of O_2_, as well as the concentration distribution of myoglobin and O_2_, is considered in domain ① (the tissue). The two cutlines along the *x* and *y* directions are marked in red while the black dot is the origin of the coordinates.

**Table 1.**
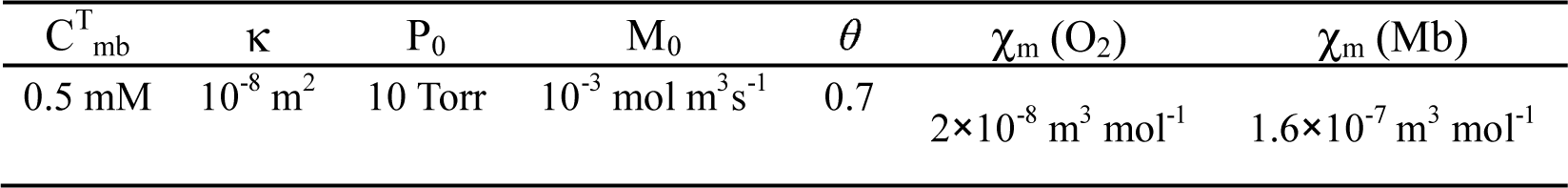
Values of selected parameters used in the model and the simulation

## 3. Results

### Interstitial flow and O_2_ transport driven by magnetic field

The distribution of magnetic flux density ***B*** with a strong gradient at the tip end is depicted in **Fig. 2** (A). At the tip, the magnetic field energy density *B*^2^/(2*μ*_0_) diminishes rapidly and provides a large energy gradient to drive the diamagnetic fluid, as shown in the inset of **Fig. 2** (B). At the center of the tip, the force on the interstitial fluid *F* = *χ_w_B*∇*B/μ*_0_ can reach 10^4^ N·m^−3^ and above as displayed in **Fig. 2** (B).

**Fig. 2.**
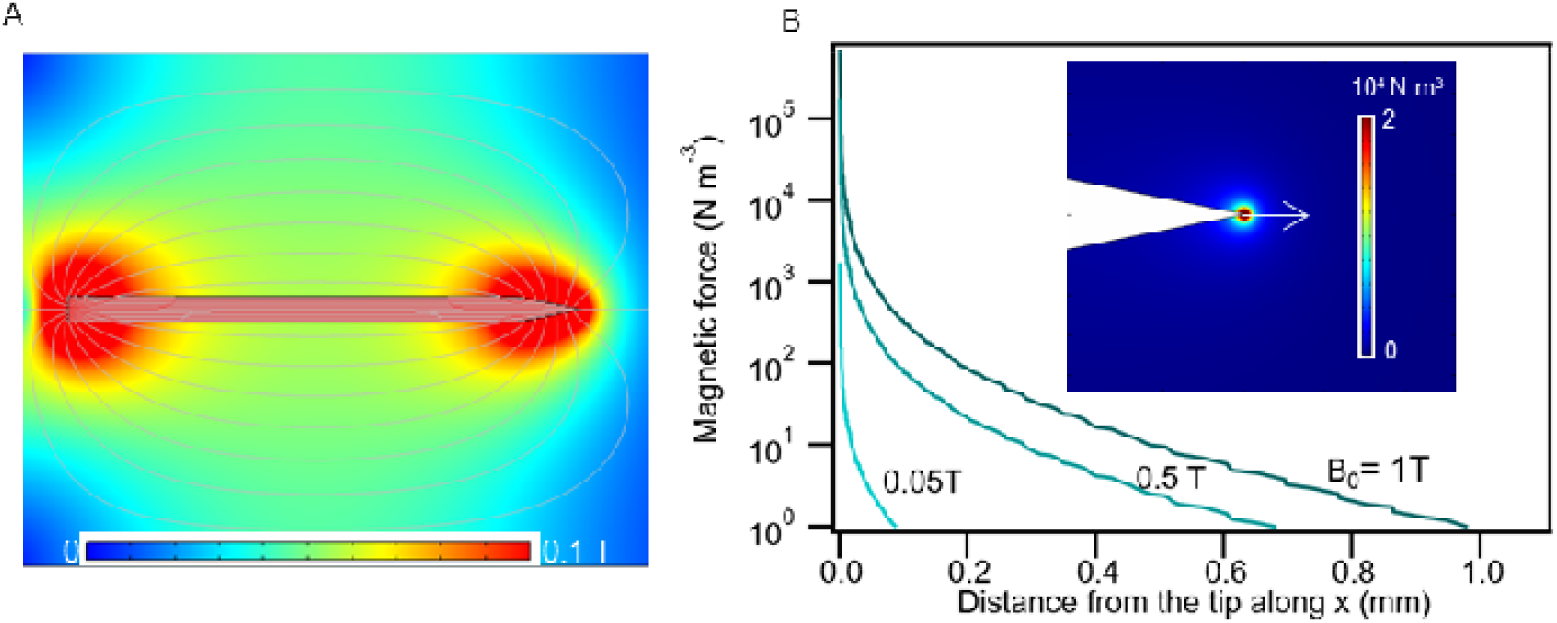
(A) Magnetic flux density of the needle. (B) Local body force exerting on the water fluid along the cutline as shown in the inset. Inset: magnetic force distribution in the vicinity of the needle tip.

The interstitial flow driven by the magnetic needle is studied inside the extracted tissue as given in **Fig 3** along the cut lines along the *x* and *y* directions (shown in **Fig. 1**). Magnetic flux of ***B*** in the tissue along the *x* and *y* directions is shown in **Fig 3** (A-B), while its magnitude depends on *B*_0_ inside needle. In the extracted tissue domain ①, the highest ***B*** is about 0.1 T for a needle with *B*_0_ = 1T. The sharp increase of a symmetric flow appears near the tip due to the high gradient magnetic field, as seen in **Fig. 3** (B). The interstitial water is expelled from the tip center and the flow rate can be above 30 μm/s at *B*_0_ = 1T. Hence, the magnetic needle inserted tissue stirs a non-uniform interstitial flow. For convenience, the average interstitial flow at *x* = 0 in tissue domain ① is used to identify the interstitial flow change with *B*_0_.

**Fig. 3.**
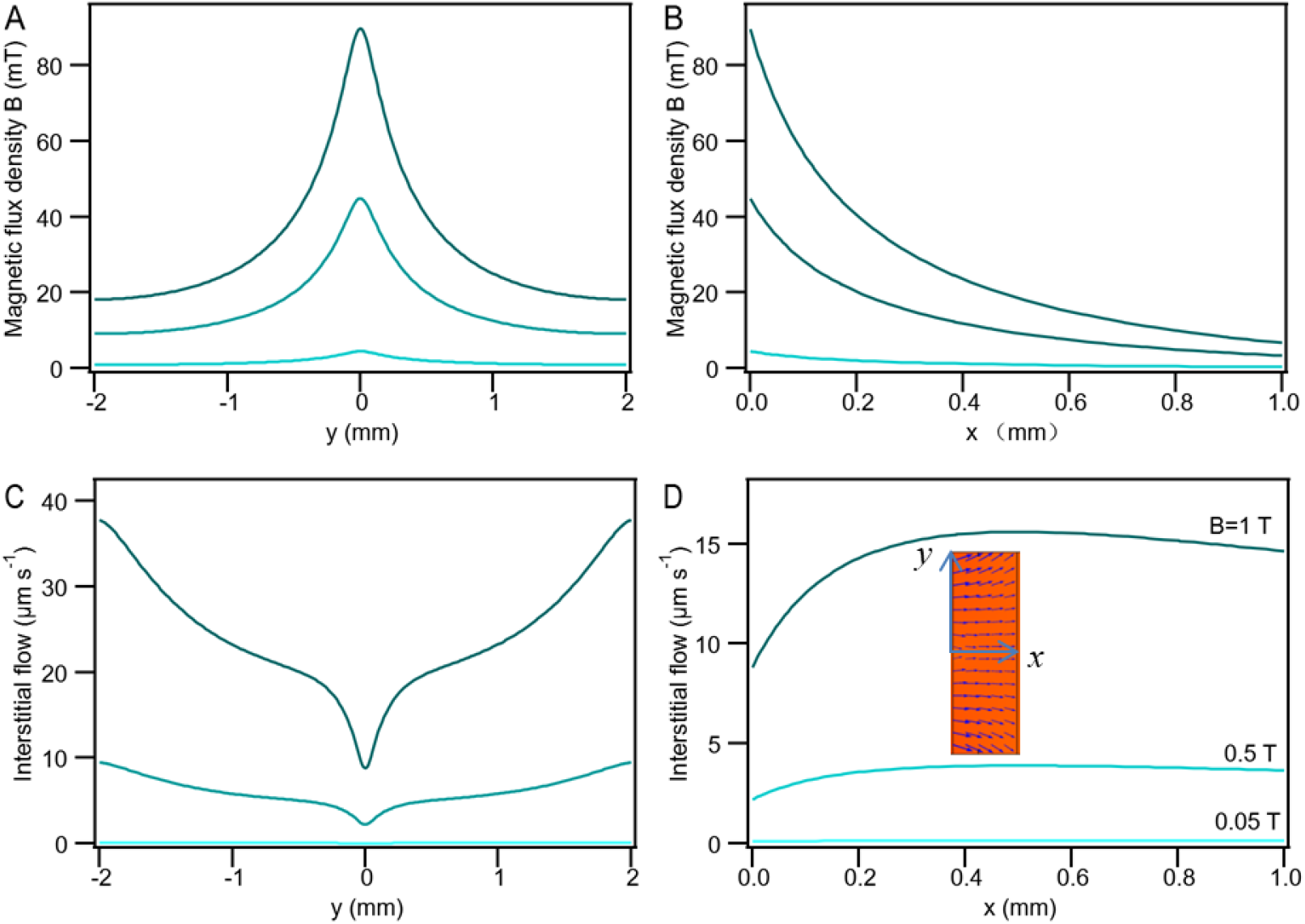
At *B*_0_ = 0.05, 0.5, and 1 T, (A-B) magnetic flux density *B* in tissue along the *y* and *x* directions; (C-D) interstitial flow along the cutlines along the *y* and *x* directions. Inset: the interstitial flow velocity distribution at *B*_0_ = 1 T.

The interstitial flow modulates the O_2_ transport through tissue. The reduced *P*_O2_ along the cut line in the *x* direction at *B*_0_ = 1T is compared to that without magnetic field, i.e., *B*_0_ = 0 in **Fig. 4** (A). As *B*_0_ increases from 0 to 1T, *P*_O2_ distributes more uniformly across tissue and more O_2_ is delivered as shown in **Fig. 4** (B). Hence, the magnetic needle accelerates the O_2_ transport by pumping a convective transport of the interstitial flow.

**Fig. 4.**
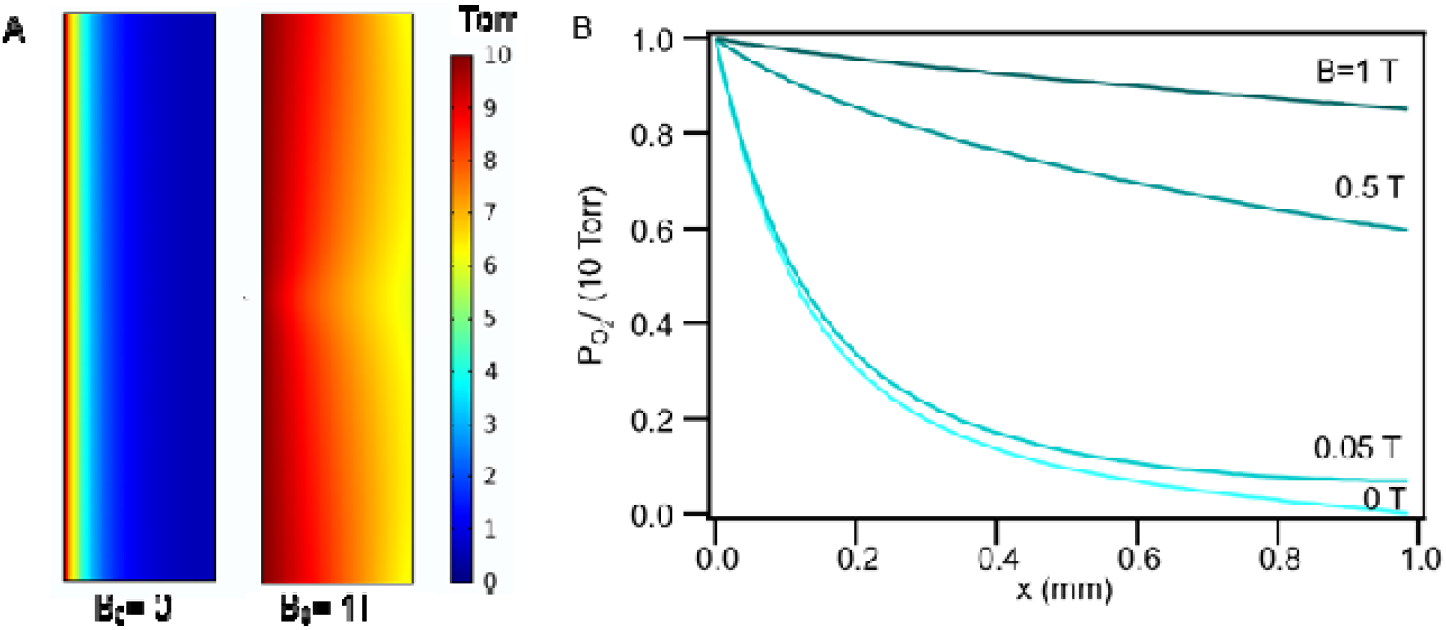
(A) *P*_O2_ profile across the tissue for the magnetic needle with the magnetic flux density *B*_0_ = 0 and 1 T, at *P*_0_ = 10 Torr. (B) O_2_ tension profile (normalized to *P*_0_) along the cutline along the *x* direction for different magnetic flux density.

To understand the active convective O_2_ transport by interstitial flow, the O_2_ input flux is used and defined as the average number of O_2_ molecules per second and area entering the tissue at *x* = 0 in domain ①. The O_2_ input flux increases dramatically with increasing *B*_0_ from 0 to 1 T, as seen in **Fig. 5** (A). Upon inserting the needle with *B*_0_ = 1T, the O_2_ input flux can reach the level with an average interstitial flow ***u*** = 22 μm s^−1^, which is nearly 50 times that without the needle intervention (***u*** = 0). To accelerate O_2_ transport by stirring the interstitial flow, *B*_0_ determines the active convective transport by interstitial flow. Even for a small *B*_0_ at 0.13 T, the contribution of the active convective O_2_ transport, through stirred interstitial flow, can be equivalent to that of a passive diffusive transport with *J*_conv_/*J*_total_ ~ 0.5. If *B*_0_ ≥ 0.5 T, the O_2_ transport through tissue is predominately carried by the interstitial flow, i.e., the active convective transport. A 2D map of the total O_2_ flux distribution inside the tissue with/without magnetic needle is displayed as the inset of **Fig. 5** (B). The average interstitial flow increases with increasing *B*_0_, as displayed in **Fig. 5** (B).

**Fig. 5.**
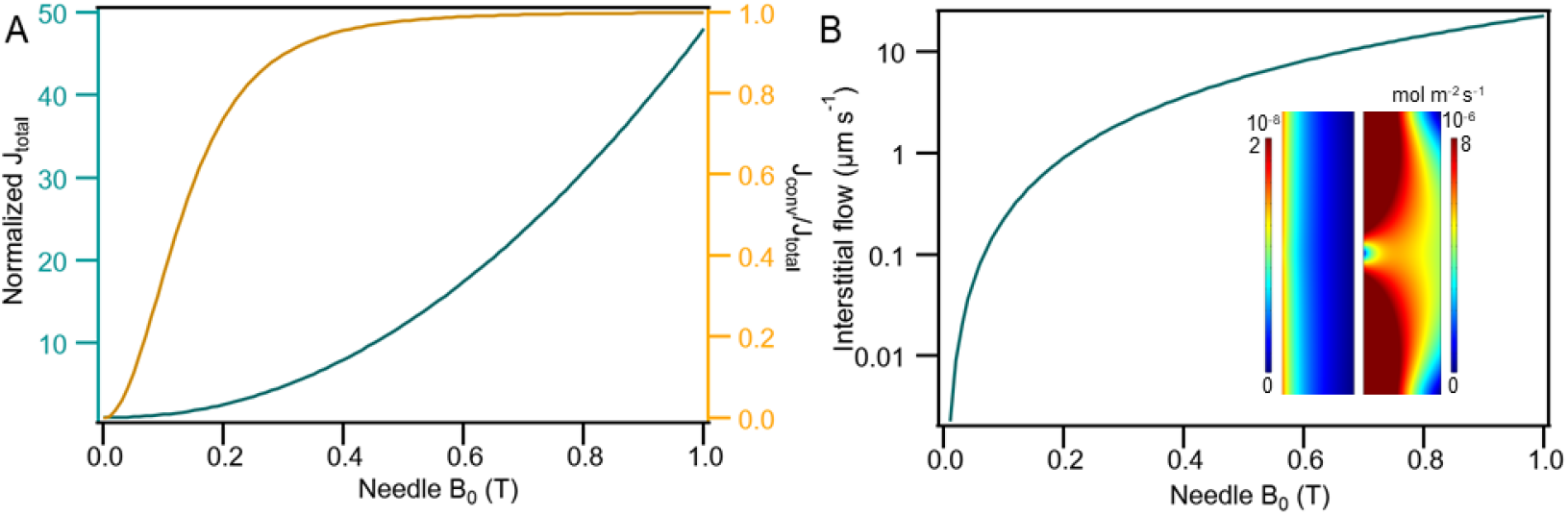
At *P0* = 10 Torr, (A) O_2_ input flux (normalized to that at *B*_0_ = 0), the ratio of *J*_conv_/*J*_total_ and (B) interstitial flow as a function of *B*_0_. The inset shows the total O_2_ flux distribution in domain ① at *B*_0_ = 0 and 1 T.

In summary, the magnetic needle provides an extra driving force to accelerate the interstitial flow that in turn triggers the active O_2_ transport inside the tissue. The active O_2_ transport in tissue is clearly correlated to *B*_0_ of the needle.

### Influence of myoglobin concentration, O_2_ tension, consumption and tissue porosity

The effect of accelerating the O_2_ transport is evaluated by varying myoglobin concentration, at *P*_0_ = 10 Torr. The normalized O_2_ input flux linearly increases with *C*_mb_, see **Fig. 6** (A), and its slope depends on *B*_0_. Therefore, the myoglobin concentration is an important factor for stimulating the O_2_ transport corresponding to stirring the interstitial flow via magnetic needle intervention. The magnetic needle near the muscle tissue with high myoglobin concentration has higher O_2_ acceleration effects compared to the muscle tissue with lower myoglobin concentration.

**Fig. 6.**
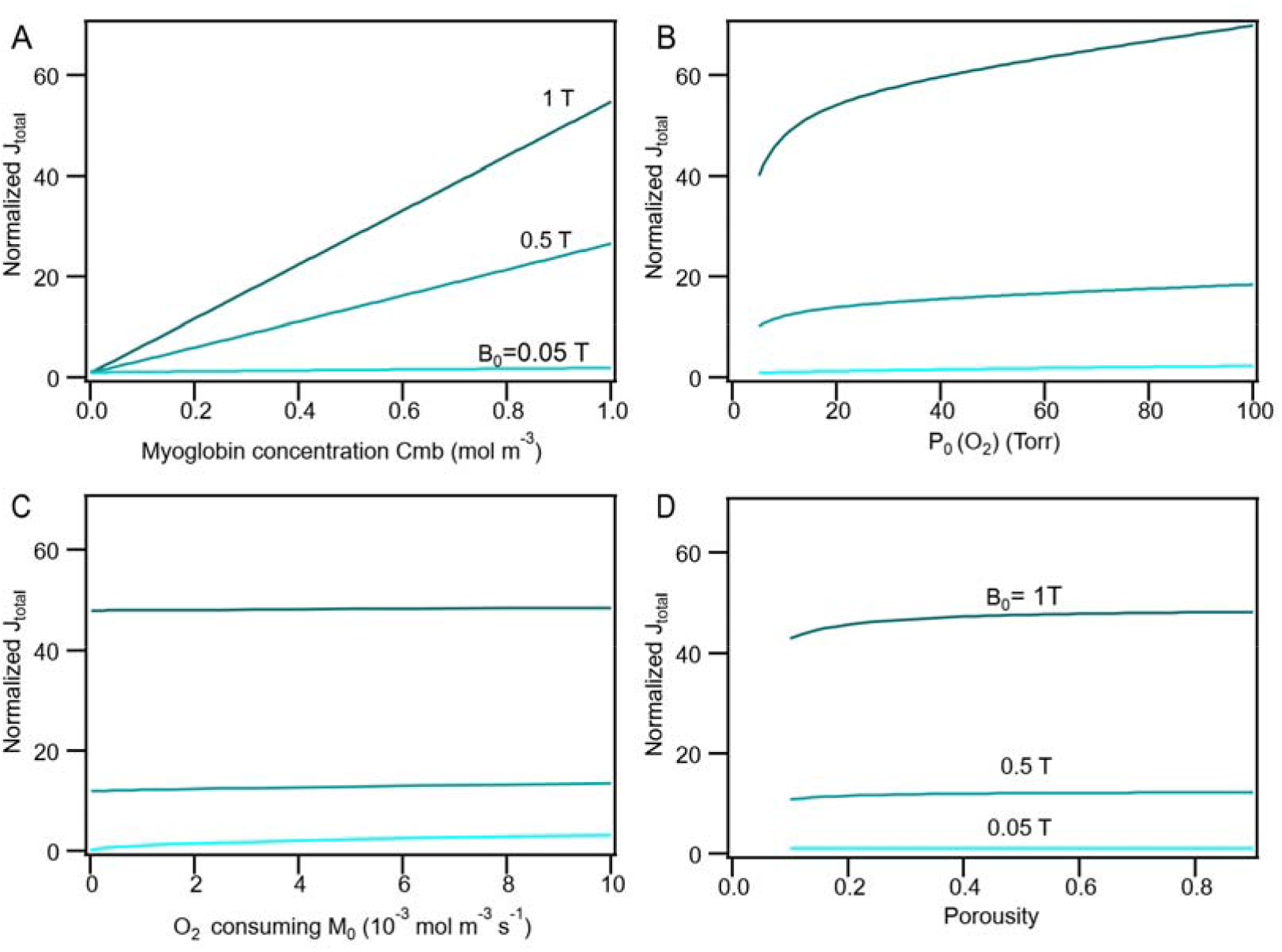
O_2_ input flux *J*_total_ at *B*_0_ = 1, 0.5, and 0.05 T as a function of (A) myoglobin concentration (normalized to that at *C*_m_b = 0), (B) O_2_ tension (normalized at to that at *P*_O2_ = 10 Torr), (**C**) O_2_ consumption rate (normalized to that at *B*_0_ = 0), and (D) porosity of the tissue (normalized to that at *B*_0_ = 0).

The O_2_ input flux as the function of *P*_O2_ in capillary is normalized to that at *B*_0_ = 0, see **Fig. 6** (B). At low *B*_0_ = 0.05 T, the enhancement of O_2_ input flux upon *P*_O2_ increase is negligible. At small *B*_0_, the enhancement also reduces the O_2_ concentration gradient in tissue with a low interstitial flow rate. Thus, the O_2_ input flux is almost constant, determined by the consumption in tissue. Only at a large value of *B*_0_ = 1 T, the enhancement of the O_2_ input flux is sensitive to *P*_O2_ in capillary. At *P*_O2_ = 100 Torr, the O_2_ input flux can reach 1.5 times that at *P*_O2_ = 10 Torr. Typical *P*_O2_ in capillary is around 40-60 Torr. Hence, the O_2_ input flux enhanced by only increasing *P*_O2_ in capillary is far less than the boosting effects of stirred interstitial flow by a magnetic needle.

The tissue porosity and O_2_ consumption rate have small effect on the O_2_ input flux as depicted in **Fig. 6** (C-D). Only at *B*_0_ = 1 T, the O_2_ input flux can increase by only 10%, as the tissue porosity changes from 0.1 to 0.9.

### Tissue permeability

The interstitial flow is correlated to the tissue permeability. Two symmetric vortices appear near the tip of the magnetic needle as the tissue permeability increases from 10^−9^ to 10^−6^ m^2^, as shown in **Fig. 7** (A). As the tissue permeability increases further, the flow ratio at the center to that at the edge of the vortex enlarges, which causes the average O_2_ input flux saturated as seen in **Fig. 7** (C). The symmetric vortices emerged in the tissue correlate to the curvature of the magnetic field and the ***B*** gradient as depicted in **Fig. 3** (A-B). The distribution of flow rate along the cut line along the *y* direction is displayed in **Fig. 7** (C). The formation of the vortex indicates the large flow rate difference in tissue with high tissue permeability. The flow pattern of these vortices is similar to the vortex current at the edge of the magnets during the electrodeposition ^47,48^.

**Fig. 7.**
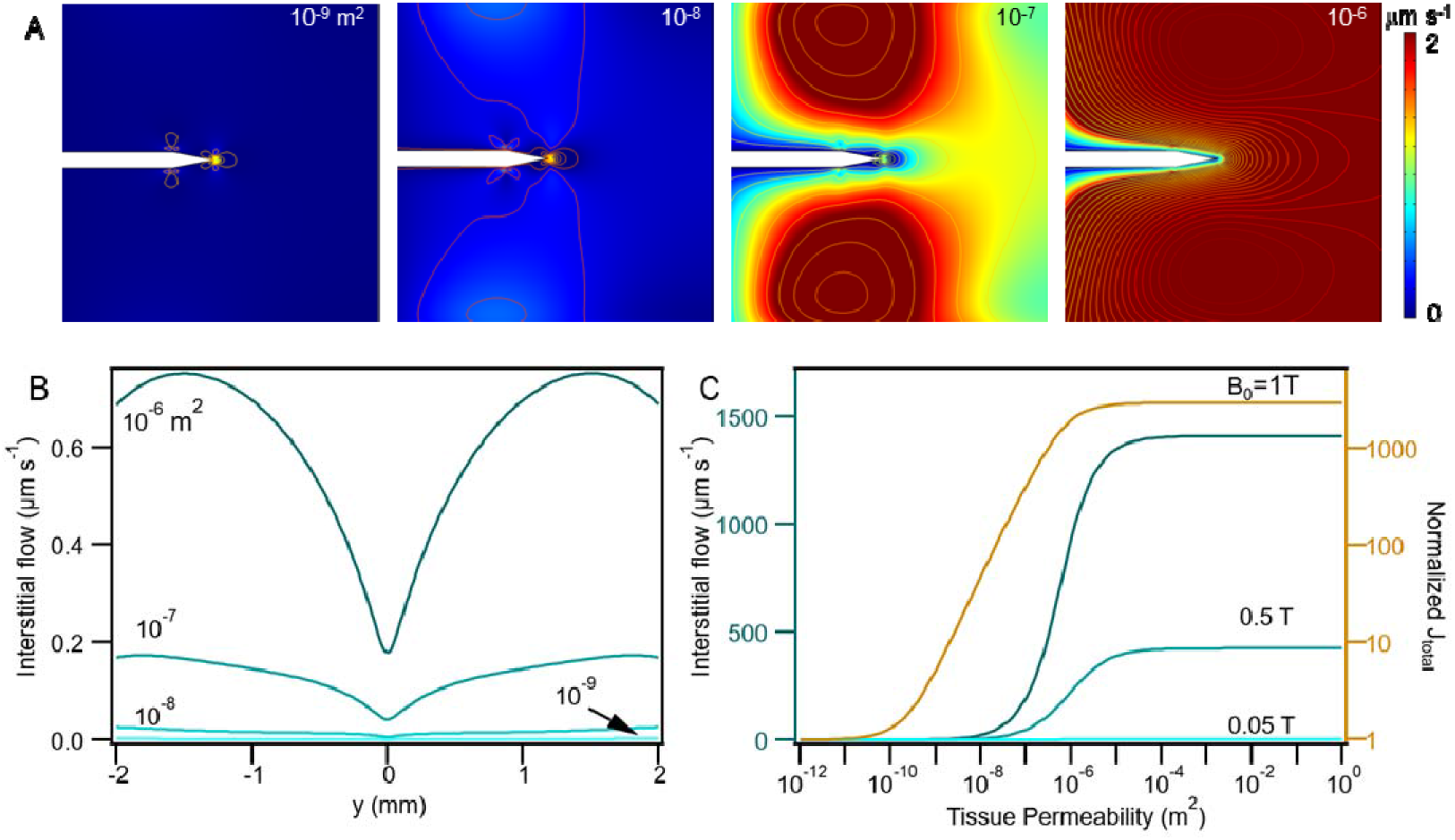
(A) Interstitial flow velocity distribution and (B) interstitial flow along the *y* direction at ***κ***=10^−6^, 10^−7^, 10^−8^, and 10^−9^. (C) Average interstitial flow rate and O_2_ input flux normalized to that at *B*_0_ = 0 as a function of specific hydraulic conductance ***κ*** for *B*_0_=1, 0.5, and 0.05 T, at *P*_0_ = 10 Torr.

### Tissue permeability surrounding the tip

Tissue permeability surrounding the needle tip is crucial in achieving fast interstitial flow rate. For different tissue permeability in domain ②, the O_2_ input flux into tissue (domain ①) with fixed *κ* = 10^−8^ m^2^ is shown in **Fig. 8**. The interstitial flow rate is mainly determined by the smallest tissue permeability in domain ① and ②. It means that, if tissue around the tip is in a health condition with low tissue permeability (10^−12^ m^2^), no interstitial flow and enhanced O_2_ transport can be observed inside edema tissue (domain ①), even domain ① and ② are very close to each other. Here, only the tip located right in a tissue with high tissue permeability can stir an interstitial flow and stimulating O_2_ transport in tissue.

**Fig. 8.**
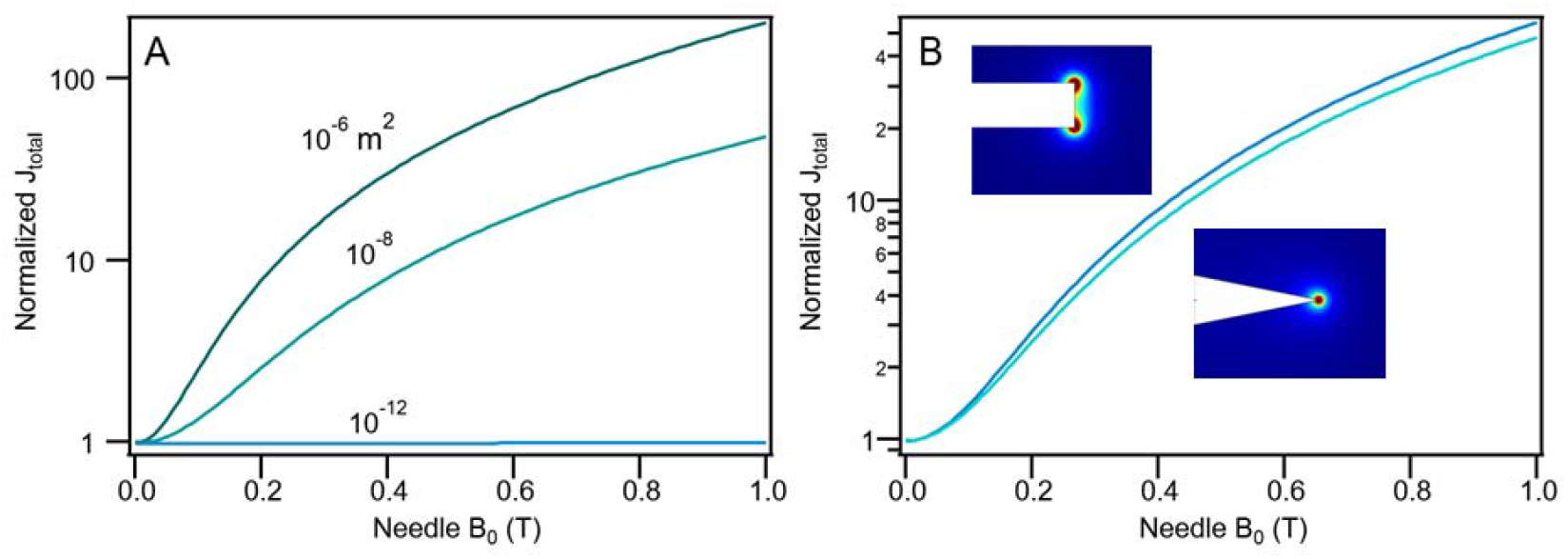
O_2_ input flux (normalized to that at *B*_0_ = 0) into the tissue (***κ*** = 10^−8^ m^2^, domain (D), (A) as a function of *B*_0_ for different tissue permeability (10^−6^, 10^−8^, and 10^−12^ m^2^) in domain ② and (B) with and without the needle tip, at *P*_0_ = 10 Torr.

Hence, the O_2_ input flux enhancement by the magnetic needle is site-specific. In addition, the needle tip shape has limited effect on the O_2_ input flux by the comparison between two needles with and without the tip, as shown in **Fig. 8** (B).

## 4. Discussion

A high gradient of magnetic field exerts a strong force on paramagnetic or diamagnetic molecules including O2, myoglobin without O_2_ loaded and water. In our case, the susceptibility of myoglobin χ(Mb) is 8×10^−8^ (at Mb = 0.5 mM) and that of O_2_ χ(O_2_) is 2.5×1010^−^ (at *P*_O2_ =10 Torr). They are 2-3 orders of magnitude lower than the water susceptibility, −9×10^−6^. Thus, the high magnetic force directly acting on myoglobin and O_2_ is limited, where its effect on O_2_ transport is also neglected. The high magnetic force was reported to enhance O_2_ dissolving in water ^39^ and cause levitation effects ^42^. Hence, the magnetic field body force on water molecules can stir a considerable interstitial flow, which promotes the active O_2_ transport via the convection process.

Hence, during the acupuncture intervention by magnetic needle, the needle provides magnetic energy to drive the movement of interstitial fluid due to the diamagnetic effect of water molecules. This interstitial flow enhances the O_2_ transport, which can pump the O_2_ input flux into the tissue (10^−8^ m^2^) with an average interstitial flow of 22 μm s^−1^ at *B*_0_ = 1 T, which is 50 times higher than that achievable without magnetic field. Also, Muscle tissue with a higher myoglobin concentration can have enlarged stimulating effects on O_2_ transport. Although, a high *P*_O2_ in capillary can enhance the O_2_ input flux to tissue, this effect is far less significance than the O_2_ transport facilitated by the magnetic needle.

The tissue permeability near the needle tip determines the interstitial flow. The needle can generate a vortex interstitial flow near the tip in the tissue of high permeability (> 10^−8^ m^2^) due to the curvature and the gradient of magnetic field near the edge of the needle tip. Moreover, the O_2_ transport enhancement is site-specific so that only the tip right located in the tissue of high permeability (> 10^−9^ m^2^, 10^−11^ 10^−15^ m^2^ m for normal tissue) has a strong effect on stimulating the O_2_ transport. However, if the pressure exerted by magnetic needle is large enough and comparable with atmospheric pressure, the tissue permeability could dramatically increase ^49^.

As discussed above, the magnetic needle can generate vortices in the interstitial flow in tissue with high tissue permeability (> 10^−9^ m^2^). Such flow redistributes O_2_ and nutrition and brings more O_2_ molecules to the damaged tissue so as to restore the mitochondria metabolism. The interstitial flow at a large rate can carry more O_2_ to the damaged tissue and restore tissue faster. Hence, the interstitial flow stirred by the magnetic needle can shift the balance state in the demand-supply cycle of O_2_ mitochondria metabolism and provide extra O_2_ for the tissue recovery.

In numerical simulation, the interstitial flow through tissue is solved in a uniform porous media. The permeability of the porous media includes the cell membrane permeability and interstitium permeability for water. For different muscle fibers, myoglobin is assumed to be uniformly distributed in the tissue. Inhomogeneities prevail in real biological systems wherein myoglobin is located inside muscle fibers and the muscle tissue is composed of different types of muscle fibers instead of a single type. Such inhomogeneities could lead to different O_2_ transport values in tissue. However, the deviations are not expected to change the conclusions reached in this work.

### Physiological mechanism of magnetic needle acupuncture

Based on the discussion above, a physiological mechanism is proposed for magnetic needle acupuncture as shown schematically in **Fig. 9**. Upon the needle insertion into the muscle tissue, the following are generated:

1. Force on tissue to produce local pressure by the magnetic needle due to the high gradient of magnetic field.
2. Pressure on tissue with a high tissue permeability (specific hydraulic conductance). If the exerted pressure is far below the atmospheric pressure, the pressure dependence of hydraulic conductance can be ignored ^49^.
3. Interstitial flow through tissue and release of the local tissue pressure. At the same time, it can

a. stimulate the active O_2_ transport through damaged tissue and
b. increase the blood volume.
4. Enhanced ATP production and O_2_ mitochondria metabolism of tissue with a facilitation of the O_2_ transport to mitochondria by 1-2 orders of magnitude.
5. Tissue repair and the microcirculation improved by an increased blood volume and active O_2_ transport.

**Fig. 9.**
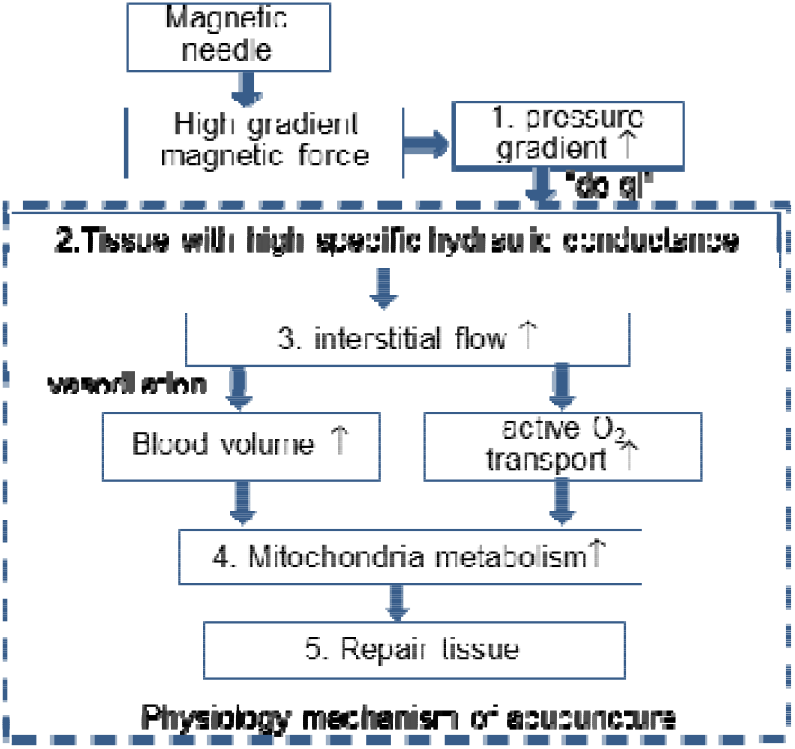
Mechanism for acupuncture.

To summarize, a magnetic needle can drive the interstitial flow to stimulate the active O_2_ transport to mitochondria to repair the local damaged tissue. This physiological mechanism can be used to account for the “de-qi” feeling as the founding of tissue with a high tissue permeability, and the blood volume increase as stirred interstitial flow and increased O_2_ demanding during acupuncture therapy.

However, the acupuncture needle could also provide a temperature gradient in the interstitial fluid to stir a flow, or a mechanical force on muscle fibers to initial the contraction to enhance the drainage of the interstitial flow. For electroacupuncture, the needle could also generate heat in the tissue so as to stir the flow, induce a current through axon, and excite the contraction of muscle fibers to facilitate O_2_ transport in the tissue. No matter what kind of needle is used, stimulating active O_2_ transport to the damaged tissue to restore the O_2_ mitochondria metabolism could be the key physiological mechanism for acupuncture.

## 5. Conclusion

The magnetic needle can drive the interstitial flow through the tissue of high tissue permeability and the active O_2_ transport in the tissue. It can also lead to pumping more than 1-2 order magnitude of the O_2_ input flux into the tissue in comparison to the O_2_ flux without magnetic effect, and restore the O_2_ mitochondria metabolism. Most importantly, it shows that the magnetic needle acupuncture may rehabilitate the local tissue metabolism by the active O_2_ transport in tissue, probably the reason why the acupuncture therapy could cure so many different diseases. We have proposed a physiological mechanism of magnetic needle acupuncture based on the active O_2_ transport to restore the O_2_ mitochondria metabolism. This mechanism can be used to explain that the “de-qi” feeling is an indicator of finding the tissue of high tissue permeability and that the observed increase of blood volume is caused by a stirred interstitial flow and increased O_2_ demanding during acupuncture therapy.

## Authors contribution

Z.L. conceived the idea. Z.L. discussed with C.W. on the physiology aspects of the idea and with S.-L.Z. on the physics aspects. Z.L. worked on model design, data analysis, and drafted the manuscript. C.W. built the COMSOL model and draft the model part. All authors contributed to the manuscript revision.

## Acknowledgements

We thank Prof. Vassilios Kapaklis, Department of Physics and Astronomy, Uppsala University for discussion of body force by magnetic needle and Shiming Zhou, Department of Physics, Tongji University, China for discussion of magnetic properties of magnetic needle.

## Physiological relevance

From the proposed acupuncture physiology, the sham point should be redefined in the acupuncture therapy. The sham point reported in the literature, such as using a blunt tip on the skin without penetration, could also pass a pressure gradient to the acupoint or the edema tissue of high tissue permeability and activate the interstitial flow and the active O_2_ transport through the tissue. Hence, the reported placebo effects of acupuncture therapy should be re-examined.

